# Multipotent tail bud cells originated with the chordate common ancestor

**DOI:** 10.64898/2026.10.25.761592

**Authors:** Tasuku Ishida, Wei-Chen Chu, Jr-Kai Yu, Yutaka Satou

## Abstract

In vertebrate embryos, the tail bud region generates the neural tube, the notochord, and somites in the posterior part of the body, including the post-anal tail. Here, we traced the fate of tail bud cells during embryogenesis in the cephalochordate, amphioxus, the earliest diverging lineage of extant chordates, and showed that in amphioxus as well, these cells contribute to the neural tube, the notochord, and somites. Our work provides direct evidence to support the hypothesis that ancestral chordates possessed tail bud cells capable of producing both neural and mesodermal derivatives, illuminating the evolutionary origin of posterior body development in chordates.

## Introduction

A post-anal tail is a synapomorphy that defines the body plan of the phylum Chordata and that tends to be associated with active locomotion (Haeckel, 1894; Kowalewski, 1867; Satoh et al., 2014). In the posterior part of vertebrate embryos, the tail bud region generates the neural tube, the notochord, and somites that constitute the post-anal tail (Cambray & Wilson, 2002; Gont et al., 1993). This tail bud region contains two multipotent progenitor populations: neuromesodermal progenitors (NMPs), which give rise to the neural tube and paraxial mesoderm (Tzouanacou et al., 2009), and midline progenitor cells (MPCs), which give rise to the notochord and floor plate of the neural tube (Row et al., 2015). Tunicates, the sister group of vertebrates, also possess a post-anal tail, although ascidians, a group of tunicates, likely lost the larval anus during evolution (Chiba et al., 2004). In the ascidian, *Ciona*, NMP-like cells contribute to the nerve cord and muscle cells near the tip of the larval tail (Ishida & Satou, 2024). Hence, the evolutionary origin of multipotent progenitor populations in the tail bud region can be traced at least as far back as the last common ancestor of tunicates and vertebrates.

Cephalochordates (amphioxus) are the sister group to Olfactores (tunicates and vertebrates). Gene expression patterns and EdU pulse-chase experiments have suggested that tail bud cells of amphioxus embryos contribute to neural, mesodermal, and endodermal cells (Gattoni et al., 2022; Schubert et al., 2001). The amphioxus tail bud region contains cells that co-express *Bra2*, an ortholog of vertebrate *Brachyury*, and *Soxb1c*, an ortholog of vertebrate *Sox1/Sox2/Sox3* (Dai et al., 2024), as do vertebrate tail bud multipotent progenitor cells, which co-express *Brachyury* and *Sox2* (Henrique et al., 2015; Row et al., 2015). These observations raise the possibility that amphioxus, like vertebrates, possesses a multipotent progenitor population in its tail bud.

## Results and Discussion

To directly test the developmental potential of the amphioxus tail bud, we traced the lineage of tail bud cells using the photoconvertible fluorescent protein, Kaede. We microinjected synthetic mRNA for *Kaede* into one- or two-cell embryos of the Florida amphioxus, *Branchiostoma floridae*, and labeled a small number of cells in the tail bud by photoconversion of Kaede from green to red fluorescence using ultraviolet (UV) light during the middle neurula stage (**Figure 1A, B**). After embryos developed to the tailbud stage, photoconverted Kaede was observed in somite cells, neural tube cells, notochord cells, and tail bud cells in most embryos examined (n = 9) (**Figure 1C–E; summarized in Figure 1F**). This observation indicates that the amphioxus tail bud region contains progenitors that contribute to the neural tube, the notochord, and somites.

**Figure 1.**
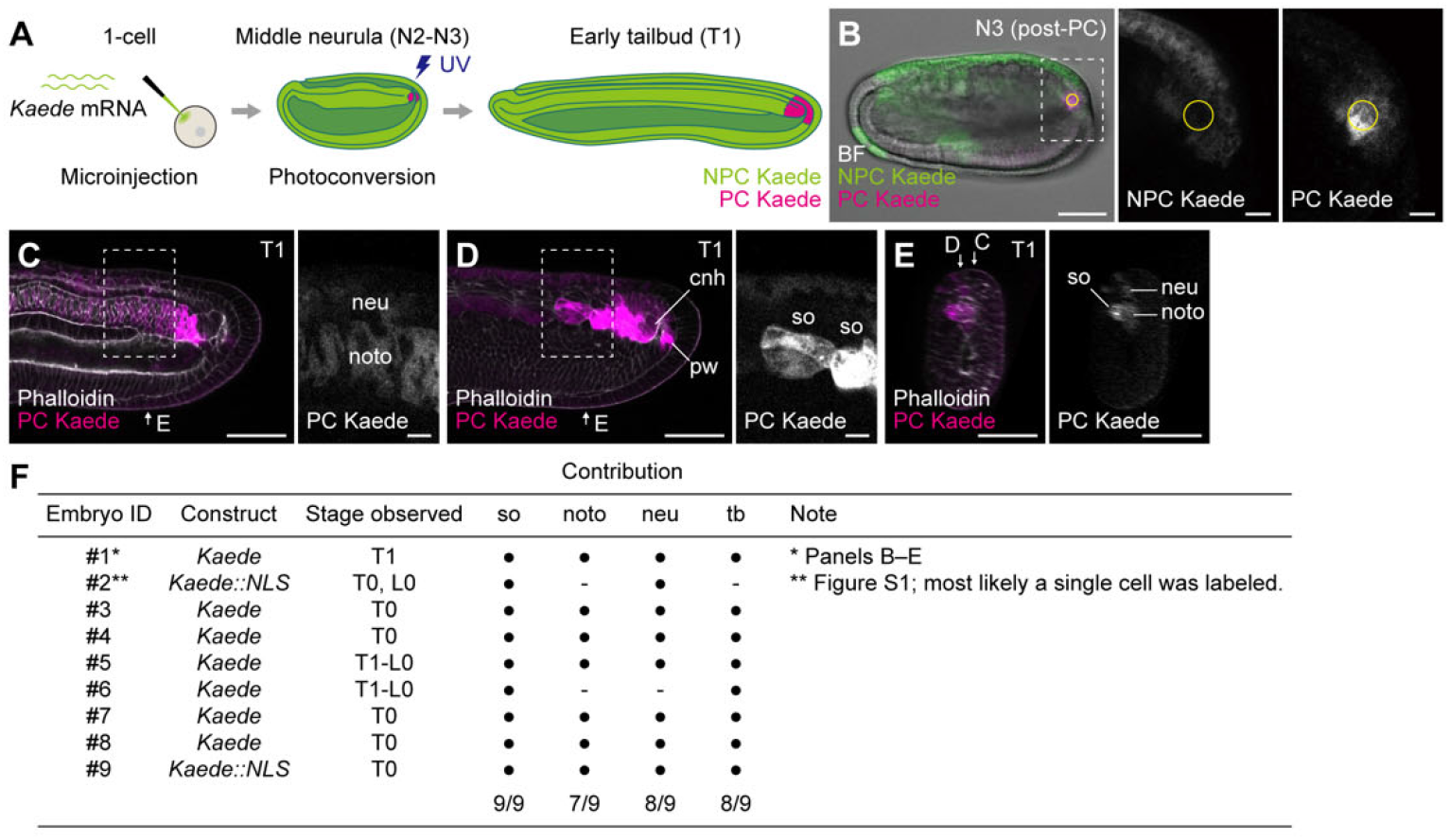
Amphioxus tail bud cells give rise to the neural tube, the notochord, and somites. (**A**) Schematic of the Kaede lineage tracing experiment shown in B–E. (**B**) Photoconversion of Kaede in the tail bud region at middle neurula stage (N3). Photographs are optical section images overlaid in pseudocolor (bright field, gray; non-photoconverted Kaede (NPC Kaede), green; photoconverted Kaede (PC Kaede), magenta). Higher-magnification views are shown on the right. Yellow circles indicate regions irradiated with UV light. (**C–E**) Photoconverted Kaede at the early tailbud stage (T1). Optical section images are overlaid in pseudocolor (PC Kaede, magenta; phalloidin, gray). C and D show two focal planes in lateral view, and E shows a cross-sectional view reconstructed from lateral-view images. Higher-magnification views are shown on the right. (**F**) Summary of photoconversion-based lineage tracing experiments. Embryos shown in B–E and in Figure S1A–E correspond to embryos 1 and 2, respectively. Each row represents a single embryo. Filled circle, PC Kaede signal detected; -, signal not detected. Scale bars: 50 μm (E, left panels of B– D); 10 µm (higher-magnification view(s) of B–D). cnh, chordoneural hinge; epi, epidermis; neu, neural tube; noto, notochord; pw, posterior wall of the neurenteric canal; so, somite; tb, tail bud; BF, bright field; post-PC, post-photoconversion.

Notably, in one of these experiments, the photoconverted region happened to contain only a very small number of labeled cells, most likely a single cell (**Figure S1A)**. In this embryo, photoconverted cells were subsequently found in both the neural tube and somites (**Figure S1B–E; summarized in Figure 1F**).

To further validate our lineage tracing results, we characterized amphioxus tail bud cells using publicly available single-cell RNA-sequencing (scRNA-seq) data from *B. floridae* (Dai et al., 2024). We used *Nanos* and *Wnt1* as markers to identify tail bud cells in the scRNA-seq data (**Figure 2A–D**) because these genes are expressed in a posterior growing region that defines the presumptive tail bud in amphioxus neurula-stage embryos (Holland et al., 2000; Wu et al., 2011). Both *in situ* hybridization and scRNA-seq data indicated that cells co-expressing *Bra2* and *Soxb1c*, which were referred to as NMP homologs in a previous study (Dai et al., 2024), constitute a very small subpopulation of tail bud cells co-expressing *Nanos* and *Wnt1* (**Figure 2B, E, F**).

**Figure 2.**
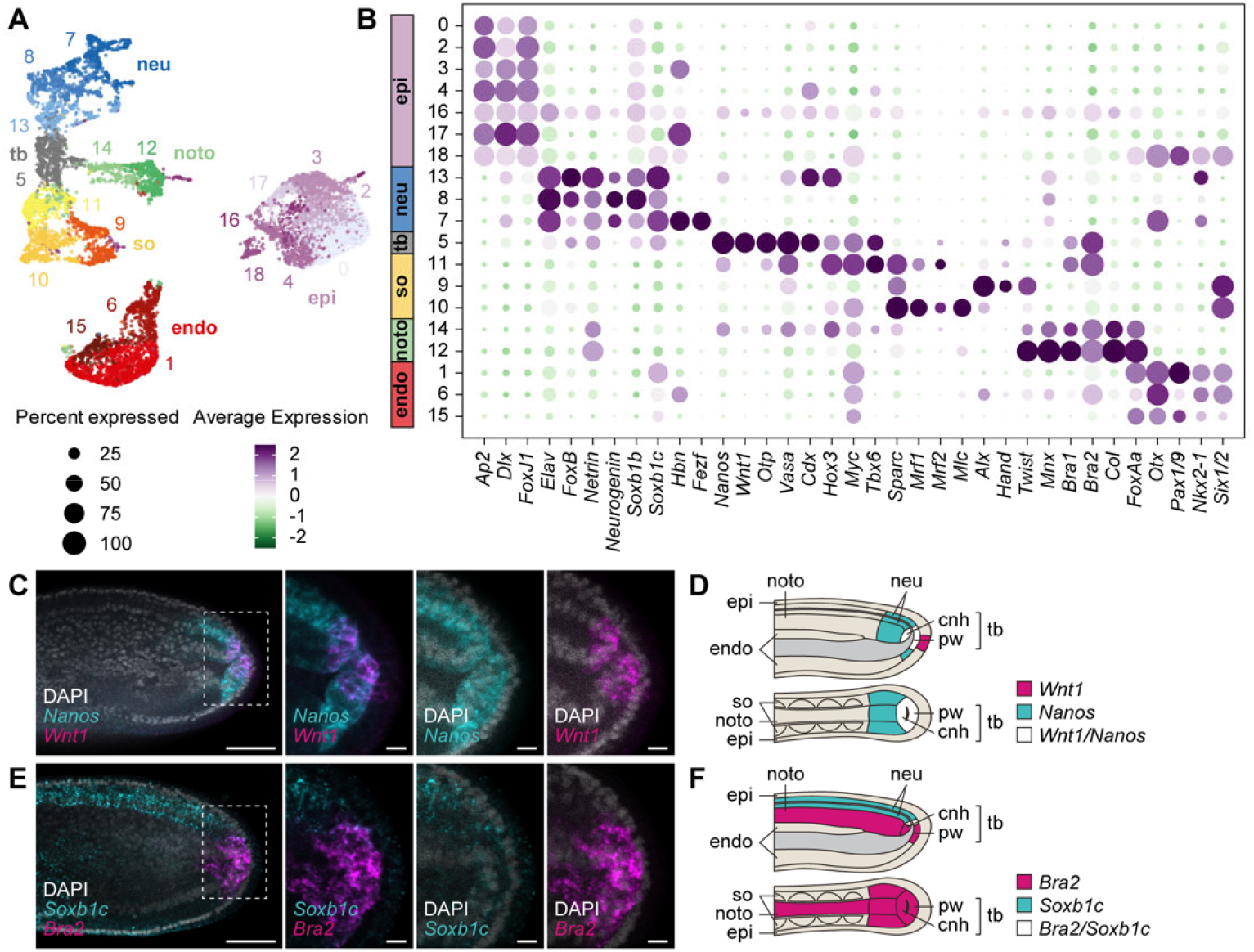
Characterization of amphioxus tail bud cells using scRNA-seq data. (**A**) UMAP plot of scRNA-seq data from *B. floridae* embryos at the middle neurula stage (N2). Clusters identified with Seurat (Hao et al., 2021) are shown in different colors with their cluster IDs. (**B**) Dot plot showing expression of markers. (**C–F**) Expression of *Nanos* and *Wnt1* (**C, D**), and of *Bra2* and *Soxb1c* (**E, F**). (**C, E**) Photographs of double fluorescence *in situ* hybridization are optical section images overlaid in pseudocolor (DAPI, gray; *Nanos* and *Soxb1c*, cyan; *Wnt1* and *Bra2*, magenta). Higher-magnification views are shown on the right. (**D, F**) Schematic representations of the anatomy of the amphioxus tail bud and expression of *Wnt1* and *Nanos* (D) and of *Bra2* and *Soxb1c* (F). Scale bars: 50 μm (left panels of C and E); 10 µm (higher-magnification views of C and E). cnh, chordoneural hinge; endo, endoderm; epi, epidermis; neu, neural tube; noto, notochord; pw, posterior wall of the neurenteric canal; so, somite; tb, tail bud.

To examine the evolutionary relationship between the amphioxus tail bud and the vertebrate tail bud region, which contains the two multipotent progenitor populations, we compared transcriptomes of amphioxus cell clusters with those of zebrafish tissue types (Lange et al., 2024) using SAMap (Tarashansky et al., 2021). This analysis revealed strong similarity of the amphioxus tail bud (cluster 5) to the zebrafish tail bud, and weak similarity to tail bud derivatives (floor plate, central nervous system, and neural plate) (**Figure 3A**). The amphioxus tail bud also showed weak similarity to zebrafish primordial germ cells (**Figure 3A**), consistent with previous findings that amphioxus tail bud cells express germline markers such as *Nanos*, *Vasa*, and *Piwi* (Wu et al., 2011; Zhang et al., 2013). Furthermore, supporting our lineage tracing results, trajectory analysis with Monocle 3 (Cao et al., 2019) revealed that tail bud cells were transcriptomically continuous with neural, notochord, and somite cells (**Figure 3B**). Thus, the amphioxus tail bud region contributes to the neural tube, the notochord, and somites, and these findings also support the hypothesis that the amphioxus tail bud region is homologous to the zebrafish tail bud region.

**Figure 3.**
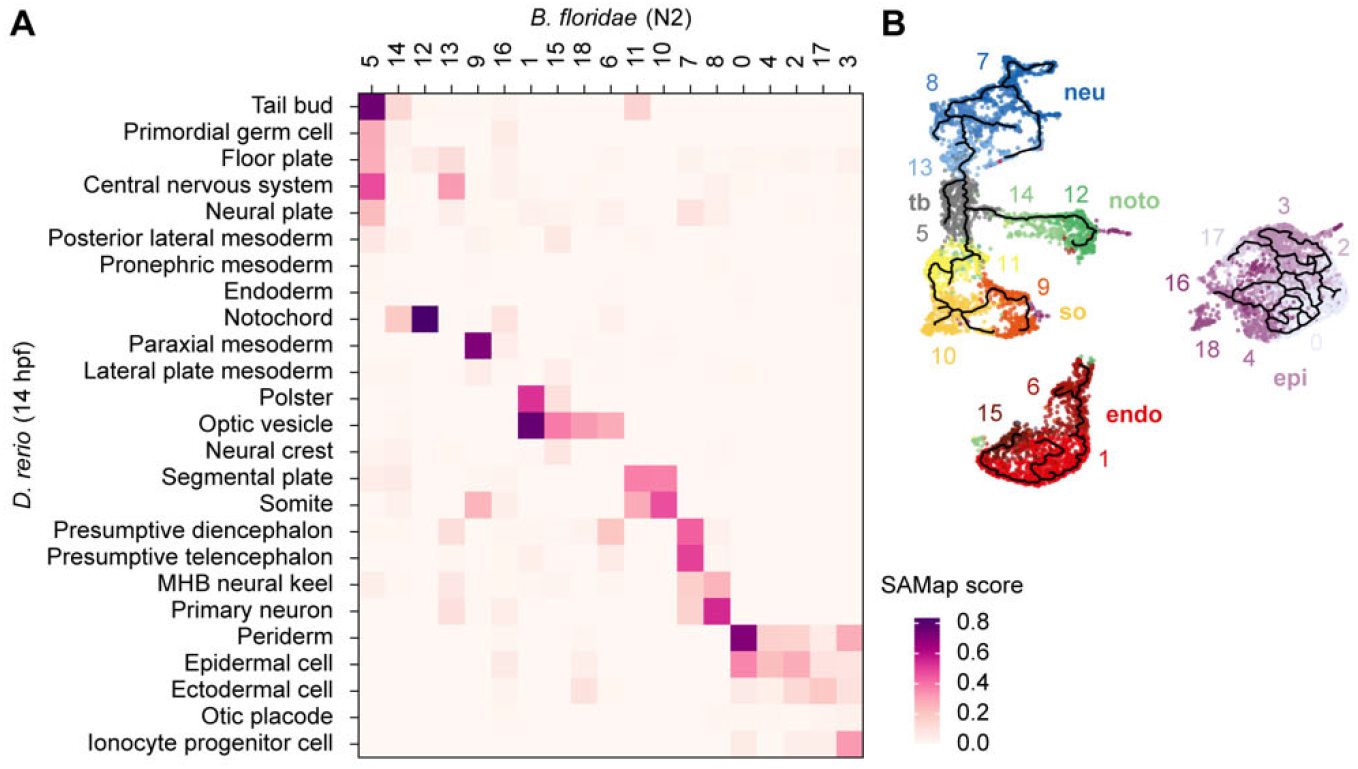
Transcriptomic comparison and trajectory analysis of amphioxus tail bud cells. (**A**) Heatmap of SAMap (Tarashansky et al., 2021) mapping scores between *B. floridae* clusters at the middle neurula stage (cluster IDs as in Figure 2A) and zebrafish tissue types at 14 hpf (Lange et al., 2024). (**B**) Transcriptomic trajectory inferred by Monocle 3 (Cao et al., 2019) (black lines), overlaid on the UMAP plot shown in Figure 2A.

The tail bud region of both amphioxus and vertebrate embryos consists of the anterior wall of the neurenteric canal (the chordoneural hinge, where the notochord and neural tube merge) and the posterior wall of the neurenteric canal (**see Figure 2D**). NMPs (located in the posterior wall) contribute to the neural tube and somites of the posterior body, whereas MPCs (located in the chordoneural hinge) give rise to the floor plate of the neural tube and notochord (Henrique et al., 2015; Row et al., 2015; Tzouanacou et al., 2009). Our lineage tracing experiments showed that amphioxus tail bud cells generate the neural tube, the notochord, and somites (**Figure 1**). These derivatives include both somites, which arise from NMPs, but not from MPCs in vertebrate embryos, and the notochord, which arises from MPCs, but not from NMPs. The amphioxus tail bud therefore generates cells that are homologous to derivatives of both vertebrate NMPs and MPCs, although the current resolution of our lineage tracing experiments cannot definitively determine whether these two potentials are combined in individual cells. Given that tunicates also possess a cell population that produces the nerve cord and muscle of the posterior body (Ishida & Satou, 2024), our results suggest that animals in all three chordate subphyla possess tail bud cells capable of producing both neural and mesodermal derivatives. It is possible that amphioxus tail bud cells may have properties of both vertebrate NMPs and MPCs. If so, the common ancestor of chordates may have possessed a tail bud multipotent cell population exhibiting combined properties of vertebrate NMPs and MPCs. We propose that evolutionary acquisition of a tail bud multipotent cell population was the crucial step that drove emergence of the post-anal tail, which is a key innovation for active locomotion in extant chordates.

## Acknowledgements

We thank Tzu-Kai Huang and personnel at the Marine Research Station and the Imaging Core Facility of ICOB for technical assistance. This research was supported by a GINPU grant from Kyoto University to TI, by grants from the Japan Society for the Promotion of Science under grant numbers 25KJ1652 to TI and 24K02032/24K21274 to YS, and by a grant from the National Science and Technology Council, Taiwan under grant number 113-2621-B-001-004-MY3 to JKY. TI was supported by the TIGP-X Pilot Program, Academia Sinica, Taipei, Taiwan.

## Author contributions

TI, JKY, and YS conceived and designed the experiments. TI and WCC performed the experiments. TI and YS analyzed the data. TI and YS wrote the original draft of the manuscript; WCC and JKY reviewed and edited the manuscript.

## Materials and Methods

### Animals and microinjection

Adult Florida amphioxus (*B*. *floridae*), derived from animals collected in Tampa Bay, Florida, United States, had been maintained in an aquarium for 12 years. Gametes were obtained by spontaneous spawning induced by temperature shifts (Holland & Li, 2021; Yong et al., 2019), and fertilization occurred spontaneously in the aquarium. Collected embryos were subsequently cultured in filtered seawater at 25–30 °C (Yong et al., 2019; Yu & Holland, 2009a), and staged as described previously (Carvalho et al., 2021).

Photoconversion-based cell-lineage tracing in *B. floridae* embryos was performed using the photoconvertible fluorescent protein, Kaede, fluorescence of which is irreversibly converted from green to red upon exposure to UV light (Ando et al., 2002; Hatta et al., 2006; Meister et al., 2022). The coding sequence of *Kaede* was inserted into the pBluescript RN3 vector (Lemaire et al., 1995). In some experiments, we used a nuclear-localized variant, Kaede::NLS, in which the sequence encoding the SV40 nuclear localization signal (PKKKRKV) was inserted upstream of the stop codon of *Kaede*. Because this variant marks nuclei and does not reveal shapes of labeled cells, it was used in combination with a membrane-tagged red fluorescent protein, mKate2::CaaX, in which the sequence encoding the C-terminal 20 amino acids of *B. floridae* GTPase HRas (NCBI accession number, XP_035696597.1; RQATKRQGKDGTSGTCCVLL), which contain the CaaX motif, was inserted upstream of the stop codon of *mKate2*. All of these constructs were inserted into pBluescript RN3 vectors, and mRNAs were synthesized *in vitro* from plasmids using the mMESSAGE mMACHINE T3 Transcription Kit (Thermo Fisher Scientific, AM1348). *Kaede* mRNA was diluted to 1 µg/µL in RNase-free water containing 1.66 µg/µL Fast Green (FUJIFILM Wako, 061-00031) and 15% glycerol (Honeywell, 15523-1L), and was injected into fertilized eggs or one blastomere of 2-cell stage embryos. *Kaede::NLS* and *mKate2::CaaX* mRNAs were mixed, each at a final concentration of 700 ng/µL, in RNase-free water containing 1.66 µg/µL Fast Green and 15% glycerol, and injected into one blastomere of 8-cell stage embryos. This restricted expression to a part of the embryo and thereby reduced the number of labeled cells in the irradiated region.

Injected embryos were incubated in filtered seawater. For photoconversion, embryos were immobilized by transfer to 2× artificial seawater (1038.4 mM sodium chloride, 22.2 mM potassium chloride, 20 mM calcium chloride dihydrate, 49 mM magnesium chloride hexahydrate, and 51 mM magnesium sulfate). Photoconversion and imaging were performed at the middle neurula stage (N2 or N3) by irradiating the target region with a 405-nm laser using an LSM880 inverted confocal microscope (Zeiss), and images were acquired using the confocal microscope with a water-immersion LD LCI Plan-Apochromat 25×/0.8 objective lens and Immersol^®^ W 2010 (Zeiss, 444969-0000-000). Photoconverted embryos were immediately washed several times with filtered normal seawater and incubated in filtered seawater. Embryos were fixed at the early tailbud (T0) to larval (L0) stages with 4% PFA-MOPS-EGTA and washed with PBST and PBSTT (0.1% Triton X-100 and 0.1% Tween20 in PBS). Embryos were blocked for 2 h in blocking solution (1% BSA in PBST) and incubated with Alexa Fluor™ 633 Phalloidin (Invitrogen, A22284) in blocking solution. Then, embryos were washed with PBST, mounted with VECTASHIELD Mounting Medium (Vector Laboratories, H-1000), and imaged with the confocal microscope.

### *In situ* hybridization

cDNA fragments of *Bra2*, *Nanos*, *Soxb1c*, and *Wnt1* were obtained by reverse transcription and PCR using the following primers: for *Bra2*, 5’-ATGTCTGAGTCTTTTCCTTG-3’ and 5’-TGGTCACAGTGTTCCTCAG-3’; for *Nanos*, 5’-TCCATGGAACGAGCATAA-3’ and 5’-CCTATGTTTTCTTACAACTTTT-3’; for *Soxb1c*, 5’-TCACATGTGACTGAGGGGAA-3’ and 5’-GGCAACAAGGCCGCCAA-3’; for *Wnt1*, 5’-CAAACGGTTACTAGCGAGAG-3’ and 5’-CGCACCTCTCAACTTAGTCG-3’. These fragments were subsequently cloned into the pBluescript II SK(-) vector using an In-Fusion HD Cloning Kit (Takara, 639649). Digoxigenin (DIG)- and dinitrophenyl (DNP)-labeled antisense RNA probes were synthesized using T7 RNA Polymerase (Invitrogen, 18033-019), DIG RNA Labeling Mix (Roche, 11277073910), and Label IT Nucleic Acid Labeling Kit, DNP (Mirus Bio, MIR 3800).

*B. floridae* embryos were fixed in 4% PFA-MOPS-EGTA fixation buffer (4% PFA, 0.1 M MOPS, 1 mM EGTA and 0.5 M NaCl) at 4 °C for more than 16 h (Yu & Holland, 2009b). Following fixation, they were washed with 80% ethanol and then PBST. For permeabilization, embryos were treated with 5 μg/mL proteinase K (Sigma-Aldrich, P2308) in PBST for 3 min at room temperature (RT). The reaction was stopped by adding 10% glycine, which was immediately removed. Subsequently, embryos were washed with 2 mg/mL glycine in PBST for 5 min, followed by two washes in PBST. Embryos were again fixed with 4% PFA in PBST for 1 h at RT, and then washed with PBST. They were pre-hybridized in hybridization buffer for at least 2 h at 60 °C. The hybridization buffer contained 50% deionized formamide, 100 μg/mL of heparin, 5× SSC, 100 μg/mL yeast tRNA, 5 mM EDTA, 1× Denhardt’s solution, and 0.1% Tween20. The hybridization buffer was replaced with fresh hybridization buffer containing a DIG- and a DNP-labeled probe, and embryos were incubated at 65 °C for at least 16 h.

After hybridization, embryos were subjected to a series of stringent washes. First, embryos were washed twice at 60 °C for 5 min in 50% deionized formamide, 5× SSC, 1% SDS, and then twice at 60 °C for 15 min in the same wash solution. Next, embryos were washed twice at 60 °C for 5 min in 50% deionized formamide, 2× SSC, 1% SDS, and then twice at RT for 15 min in the same wash solution. Then, they were washed twice with 2× SSC, 0.1% Tween 20. Embryos were treated for 30 min at 37 °C with 20 μg/mL RNase A in 2× SSC, 0.1% Tween 20. Embryos were washed once in 2× SSC, 0.1% Tween 20 without incubation, twice at room temperature for 20 min in 2× SSC, 0.1% Tween 20, once in 0.2× SSC, 0.1% Tween 20 for 20 min, and once for 5 min in PBST containing 2 mg/mL bovine serum albumin. For two-color fluorescent detection, embryos were treated for 30 min in 100 mM NaN_3_ in 0.1 M Tris-HCl (pH 7.5), 0.15 M NaCl, 0.05% Triton-X 100 (TBSTx), and then washed four times for 5 min in TBSTx. Embryos were blocked at room temperature for 30 min with 0.5% blocking solution in TBSTx, and then exposed to rabbit anti-dinitrophenol (Invitrogen, A6430; 1/2,000 dilution) in Can Get Signal Immunostain Immunoreaction Enhancer Solution B (Toyobo, NKB-601) at room temperature for 1 h. After washing with TBSTx, embryos were incubated with an HRP-conjugated secondary antibody (Thermo Fisher, B40922) at room temperature for 1 h. Embryos were washed four times with TBSTx. Then, they were stained with Alexa Fluor 488 tyramide reagent (Thermo Fisher, B40953). Embryos were washed with TBSTx, then incubated for 30 min in TBSTx, and then washed with TBSTx. For the second color, they were blocked at room temperature for 30 min with 0.5% blocking solution in TBSTx, and then exposed to mouse anti-DIG (Roche, 11333062910; 1/100 dilution) in Can Get Signal Immunostain Immunoreaction Enhancer Solution A at room temperature for 1 h. After washing with TBSTx, embryos were incubated with an HRP-conjugated secondary antibody (Thermo Fisher Scientific, B40961) at room temperature for 1 h. Embryos were washed four times with TBSTx. Then, they were stained with Alexa Fluor 555 tyramide reagent (Thermo Fisher, B40955). Embryos were washed with TBSTx and counterstained with 5 ng/µL DAPI (4’,6-Diamidino-2-phenylindole Dihydrochloride; Nacalai Tesque Inc., 11034-56) in TBSTx. Then, they were mounted with VECTASHIELD Mounting Medium and imaged by microscopy.

### Data analysis

Previously published single-cell transcriptome data for *B. floridae* (Dai et al., 2024) were obtained from the Chinese National Genomics Data Center (accession number: CRR767589) and mapped to the genome sequence (Simakov et al., 2020) using CeleScope (Singleron, version 1.7.2). The genome sequence and accompanying gene annotation were obtained from the NCBI database (accession number: GCF_000003815.2). Mapped data were analyzed with Seurat (version 5.2.1) (Hao et al., 2021). Seurat objects were created using the ‘CreateSeuratObject’ function with the parameters ‘min.features=200’ and ‘min.cells=3’ (resulting in 9,935 cells). Data were normalized using the ‘NormalizeData’ function with default parameters, and variable features were identified using the ‘FindVariableFeatures’ function with default parameters. After scaling data with the ‘ScaleData’ function, principal component analysis was performed with the ‘RunPCA’ function. To cluster cells, the ‘FindNeighbors’ function was used with the parameter ‘reduction = “pca”’ and the ‘FindClusters’ function was used with default parameters. Data were visualized with the ‘RunUMAP’ function with the parameter ‘dims = 1:50’. After the first clustering, clusters corresponding to neural, somite, notochord, and tail bud cells were selected, and clustering was performed again using the ‘cluster_cells’ function. Using this clustering information, trajectory analysis was performed with Monocle 3 (version 1.37) (Cao et al., 2019) using the ‘learn_graph’ function with the parameter ‘use_partition = T’. All commands used are shown in Supplementary Text S1.

For cross-species transcriptome comparison, processed single-cell transcriptome data for zebrafish at 14 hpf were downloaded from ZebraHub (https://zebrahub.sf.czbiohub.org/; zf_atlas_14hpf_v4_release.h5ad) (Lange et al., 2024). Cross-species comparison was performed using SAMap (version 3.0.1) (Tarashansky et al., 2021), which calculated cell type transcriptomic alignment scores between amphioxus and zebrafish. Mapping scores were visualized as heatmaps using R (version 4.5.1).

## Data availability

All data generated during this study are included in this published article.

## Code Availability

R-markdowns for analyses using Seurat and monocle3 are included in Supplementary Text S1.

---

**Figure S1.**
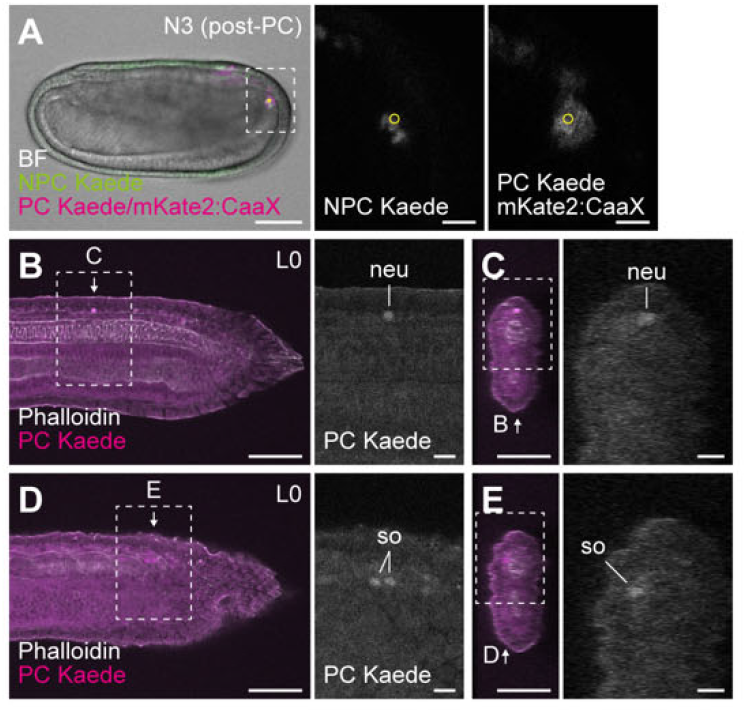
The amphioxus tail bud likely contains multipotent progenitor cells. (**A**) Photoconversion of Kaede::NLS in the tail bud region at the middle neurula stage (N3). Photographs are optical section images overlaid in pseudocolor (bright field, gray; NPC Kaede::NLS, green; PC Kaede::NLS and mKate2::CaaX, magenta). Higher-magnification views are shown on the right. Yellow circles indicate regions irradiated with UV light. (**B–E**) Photoconverted Kaede::NLS at the early larval stage (L0). Optical section images are overlaid in pseudocolor (PC Kaede::NLS, magenta; phalloidin, gray). B and D show two different focal planes in lateral view, and C and E show cross-sectional views reconstructed from the lateral-view images. Higher-magnification views are shown on the right. Scale bars: 50 μm (left panels of A–E); 10 µm (higher-magnification view(s) of A–E). neu, neural tube; so, somite. BF, bright field; post-PC, post- photoconversion.

## Supplementary Text S1. R-markdowns in analyses of transcriptome data

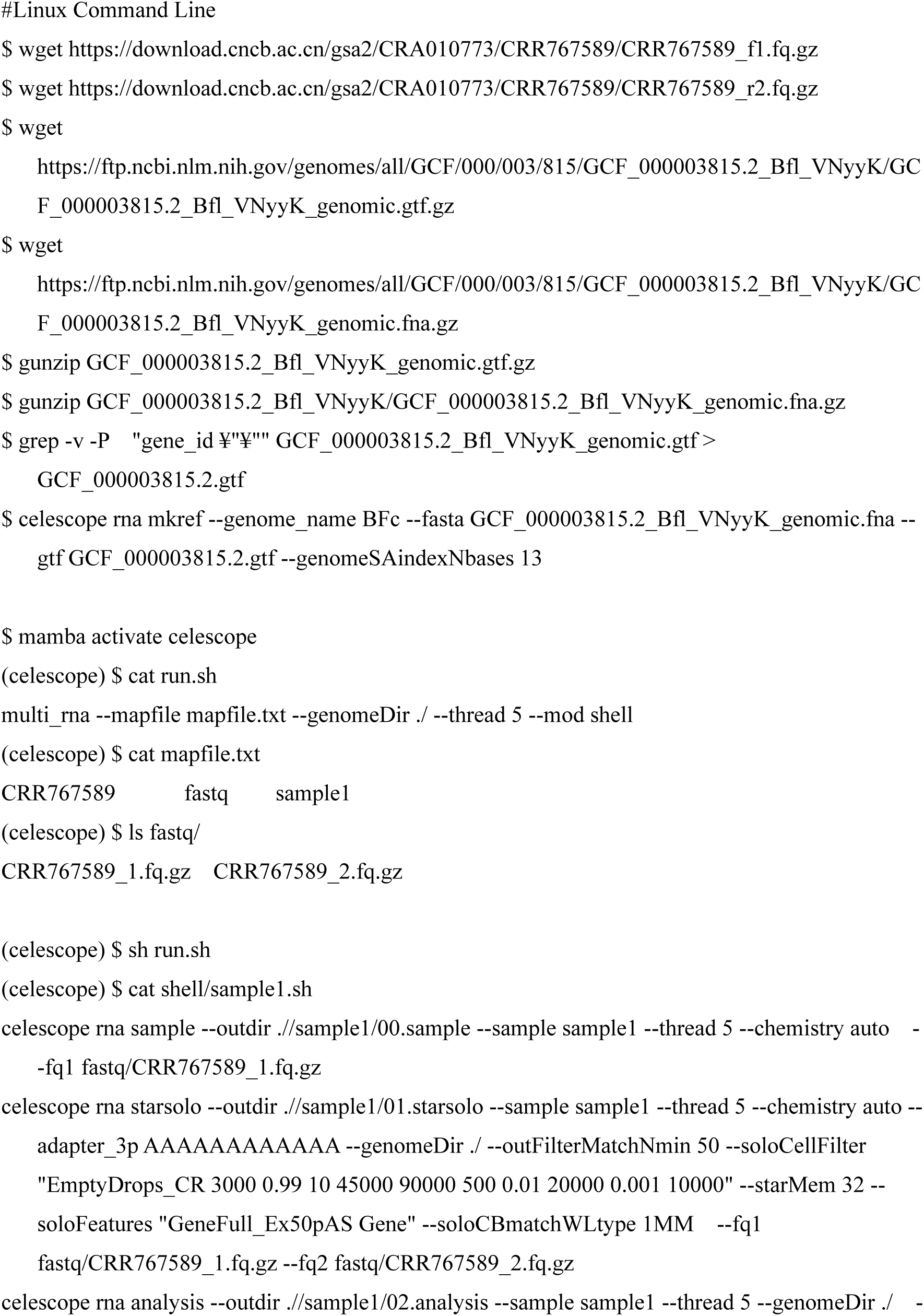
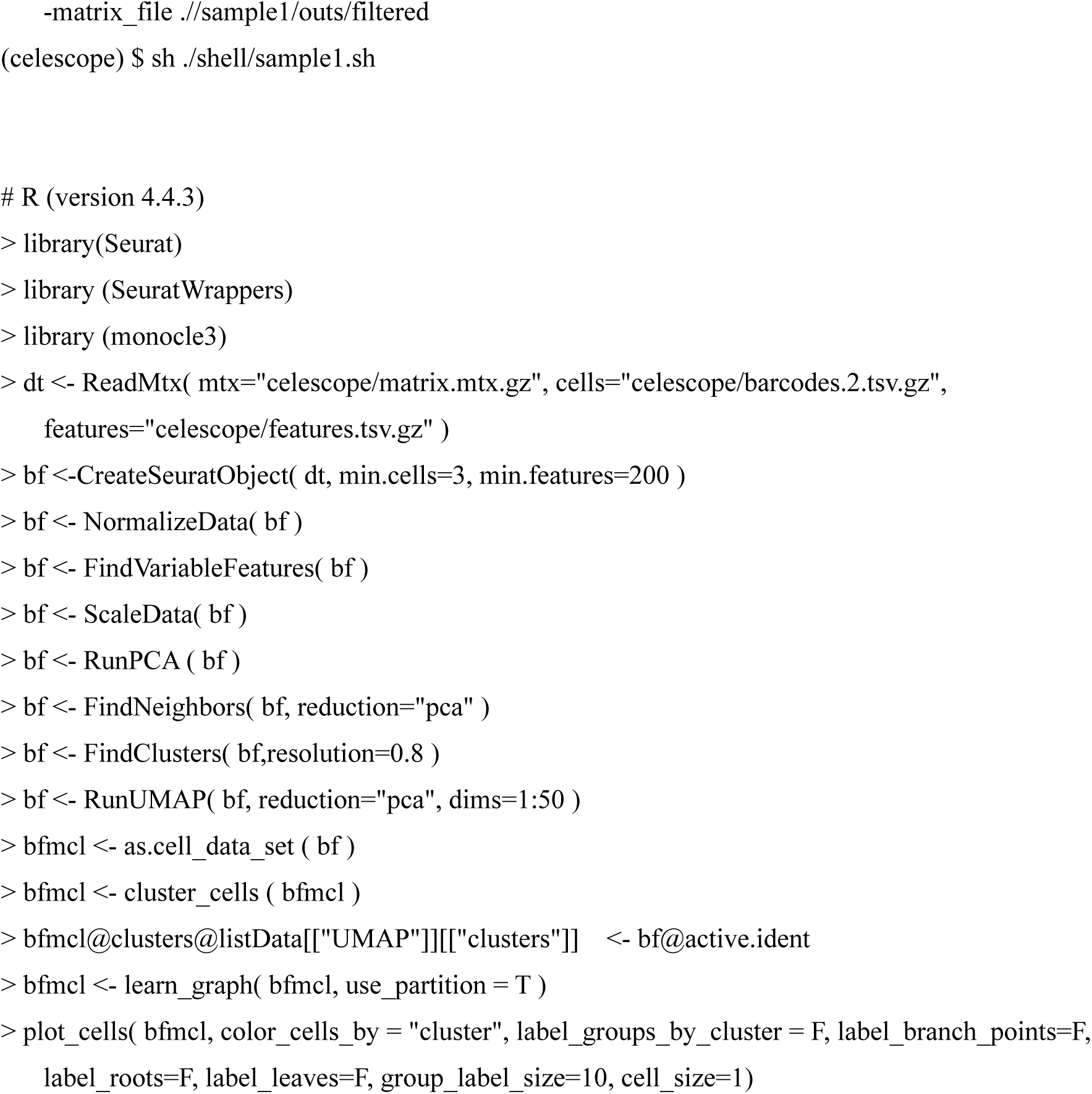

